# Acute expression of human APOBEC3B in mice causes lethality associated with RNA editing

**DOI:** 10.1101/2022.06.01.494353

**Authors:** Alicia Alonso de la Vega, Nuri Alpay Temiz, Rafail Tasakis, Kalman Somogyi, Eli Reuveni, Uri Ben-David, Albrecht Stenzinger, Tanja Poth, Nina Papavasiliou, Reuben S. Harris, Rocio Sotillo

**Affiliations:** Division of Molecular Thoracic Oncology, German Cancer Research Center (DKFZ), Im Neuenheimer Feld 280, 69120 Heidelberg, Germany; Translational Lung Research Center Heidelberg (TRLC), German Center for Lung Research (DZL); Health informatics Institute, University of Minnesota, Minneapolis, USA, 55455; Division of Immune Diversity, German Cancer Research Center (DKFZ), Im Neuenheimer Feld 280, 69120 Heidelberg, Germany; Department of Human Molecular Genetics and Biochemistry, Faculty of Medicine, Tel Aviv University, Tel Aviv, Israel.; Institute of Pathology, University Hospital Heidelberg, Heidelberg, Germany; Department of Biochemistry and Structural Biology, University of Texas Health San Antonio, San Antonio, Texas, USA, 78229; Howard Hughes Medical Institute, University of Texas Health San Antonio, San Antonio, Texas, USA, 78229

## Abstract

RNA editing has been described to promote heterogeneity leading to the development of multiple disorders including cancer. The cytosine deaminase APOBEC3B is known to fuel tumor evolution through DNA mutagenesis, but whether it may also function as an RNA editing enzyme has not been studied. Here, we engineered a novel doxycycline-inducible mouse model of human *APOBEC3B*-overexpression to understand the impact of this enzyme in tissue homeostasis and address a potential role in C-to-U RNA editing. Elevated and sustained levels of APOBEC3B led to rapid alteration of cellular fitness, major organ dysfunction, and ultimately lethality in mice. Importantly, extensive analyses of RNA-sequencing and WES from mouse tissues expressing high APOBEC3B levels reveal frequent UCC-to-UUC RNA editing events mainly localized in a specific hotspot. This work identifies, for the first time, a new function for APOBEC3B in RNA editing and presents a valuable preclinical tool to understand the emerging role of APOBEC3B as a potent driver of cancer and other diseases.

## Introduction

RNA editing is emerging at the forefront of epitranscriptomics having a fundamental role in multiple human diseases, including cancer (Baysal et al., 2017; Kung et al., 2018; Kurkowiak et al., 2021; Rayon-Estrada et al., 2015). This mechanism generates changes at the RNA level regulating genetic plasticity and resulting in protein diversity. RNA editing is a co- or post-transcriptional modification that most frequently involves the conversion of cytosine to uracil (C-to-U) or adenosine to inosine (A to I) by APOBEC or ADAR family of deaminases, respectively (Lerner et al., 2018). Although A-to-I editing has been reported extensively in thousands of positions in different species, C-to-U editing has been investigated to lesser extents. The first C-to-U editing was described in apolipoprotein B (*ApoB*) mRNA which results in a stop codon (UAA) and the generation of the truncated protein ApoB48 (Powell et al., 1987). *ApoB* editing is associated with the metabolism of lipoproteins and impaired editing increases the risk of cardiovascular disease (Powell-Braxton et al., 1998; Xie et al., 2007). To date, only a few studies have directly connected C-to-U conversion to biological processes including cancer (Alqassim et al., 2021; Cappione et al., 1997; Mukhopadhyay et al., 2002; Yamanaka et al., 1997).

RNA C-to-U editing activity appears to be restricted to a few APOBEC members. APOBEC1, the first member identified as an RNA editing enzyme is responsible for the editing of *ApoB* mRNA (Teng et al., 1993). Overexpression of Apobec1 in mice and rabbits lead to hepatocellular carcinomas showing hyperediting of multiple cytosines at different sites from the canonical one on the *ApoB* mRNA (Yamanaka et al., 1995), suggesting that high levels of APOBEC1 results in the loss of editing fidelity. In addition, under hypoxic conditions and interferon stimulation, the upregulation of APOBEC3A (A3A) has been associated with increased RNA editing in macrophages (Sharma et al., 2015). Human tumors expressing high levels of A3A also exhibited a large proportion of edited genes (Jalili et al., 2020). Lastly, APOBEC3G (A3G) has shown editing activity in HEK293T and lymphocyte cells (Sharma et al., 2019). These deaminases are promiscuous when selecting their substrate as they also have the ability to deaminate single-stranded DNA cytosines (Alqassim et al., 2021; Barka et al., 2022; Harris et al., 2002; Petersen-Mahrt & Neuberger, 2003; Saraconi et al., 2014; Sharma et al., 2015). In comparison, other family members such as APOBEC3B (A3B), show a nucleic acid substrate preference, 5’-TCW (W = A or T), that so far has only been described to function at the DNA level (Burns, Lackey, et al., 2013).

APOBEC expression leads to the accumulation of mutations as well as DNA damage, which may compromise cellular integrity (Burns, Lackey, et al., 2013; Burns, Temiz, et al., 2013). Indeed, clonal DNA sequencing revealed that A3A and A3B may be expressed in episodic bursts suggesting that continuous expression of these deaminases may be toxic for cancer cells (Petljak et al., 2019). In line with these results, the presence of APOBEC mutational signatures in human tumors does not always correlate with expression levels of A3A/B (Jalili et al., 2020; Petljak et al., 2019). Unlike DNA mutations, RNA editing is a dynamic process which does not leave a permanent footprint and RNA edits disappear right after the responsible enzymes are no longer expressed. Indeed, a recent study indicated that A3A activity can be detected by monitoring RNA editing hotspots (Jalili et al., 2020). Although several studies have addressed the relevance of RNA editing in cancer and other diseases (Kung et al., 2018; Kurkowiak et al., 2021) the complexity of capturing the labile editing scenario could have masked editing activity by other APOBEC members. Therefore, whether A3B represents an epigenetic threat to RNA and cellular integrity has yet to be investigated.

Here, we report the development of a novel doxycycline-inducible mouse model expressing the human *A3B*. Upon doxycycline administration, high and sustained A3B levels are achieved in different tissues which trigger early lethality. Whole exome and transcriptome analysis of *A3B*-expressing tissues identified hundreds of *A3B*-induced RNA editing events with 5’-U<u>CC</u> as a preferred sequence context. Some of the *A3B*-edited positions appear to be hotspots that occur in the same RNA substrate in several distinct tissues in different animals. Importantly, continuous expression of *A3B* is required for detecting RNA editing, since complete silencing of the *A3B* transgene results in the lack of editing. Together our results identify a new function for *A3B* in RNA editing, which expands our understanding of the broader APOBEC family of editing enzymes and provides new opportunities to investigate the consequences of *A3B* expression in cancer and other diseases.

## Results

### Generation of an inducible mouse model for human APOBEC3B

To investigate the consequences of A3B activity *in vivo*, human *APOBEC3B* fused to turboGFP (hereafter *A3B*) was introduced into the *ColA1* locus of KH2 mouse embryonic stem cells (Beard et al., 2006) (Figure 1A). In this system, *A3B* expression is placed under the control of a tetracycline-inducible operator (*tetO*) sequence (Figure 1A). Mice containing *A3B* were crossed with a constitutive *CAGs-rtTA3* transgene (Figure S1A) that results in robust expression of the rtTA transactivator in most adult tissues (Dow et al., 2014). Adult 4-week-old *TetO-A3B- tGFP/CAGs-rtTA3* (*A3B*) mice were feed ab libitum with doxycycline (dox)-containing diet to systematically overexpress the *A3B* transgene. 10 days after daily doxycycline exposure, high levels of tGFP were predominantly found in liver and pancreatic tissue (Figure 1B and S1B). We found human A3B stains most intensely in liver and pancreatic tissues and the enzyme localizes predominantly to the nuclear compartment (Figure 1C and S1C). Protein expression levels by immunoblotting also show A3B highly expressed in liver and pancreas, with intermediate levels in the intestine, while a minimal level of A3B protein was detected in lung and spleen (Figure 1B, C, D). We then sought to demonstrate if our *A3B* transgene retain its deaminase activity. Single-stranded DNA C-to-U activity assay with soluble protein extracts from *A3B*-expressing tissues demonstrated that A3B is functionally active (Figure 1E). To compare A3B expression levels in mice to those in humans, mRNA expression from livers, pancreas and lungs from *TetO-A3B/CAGs-rtTA3* mice was normalized to the housekeeping gene encoding TATA-binding protein (TBP). Lung tissues showed A3B expression within the range observed across human cancers, while A3B expression in liver and pancreas was comparable to human tumors exhibiting the highest A3B levels which have been associated with poor survival in patients (Law et al., 2016; Wang et al., 2018; Zhang et al., 2021) (Figure S1D).

**Figure 1.**
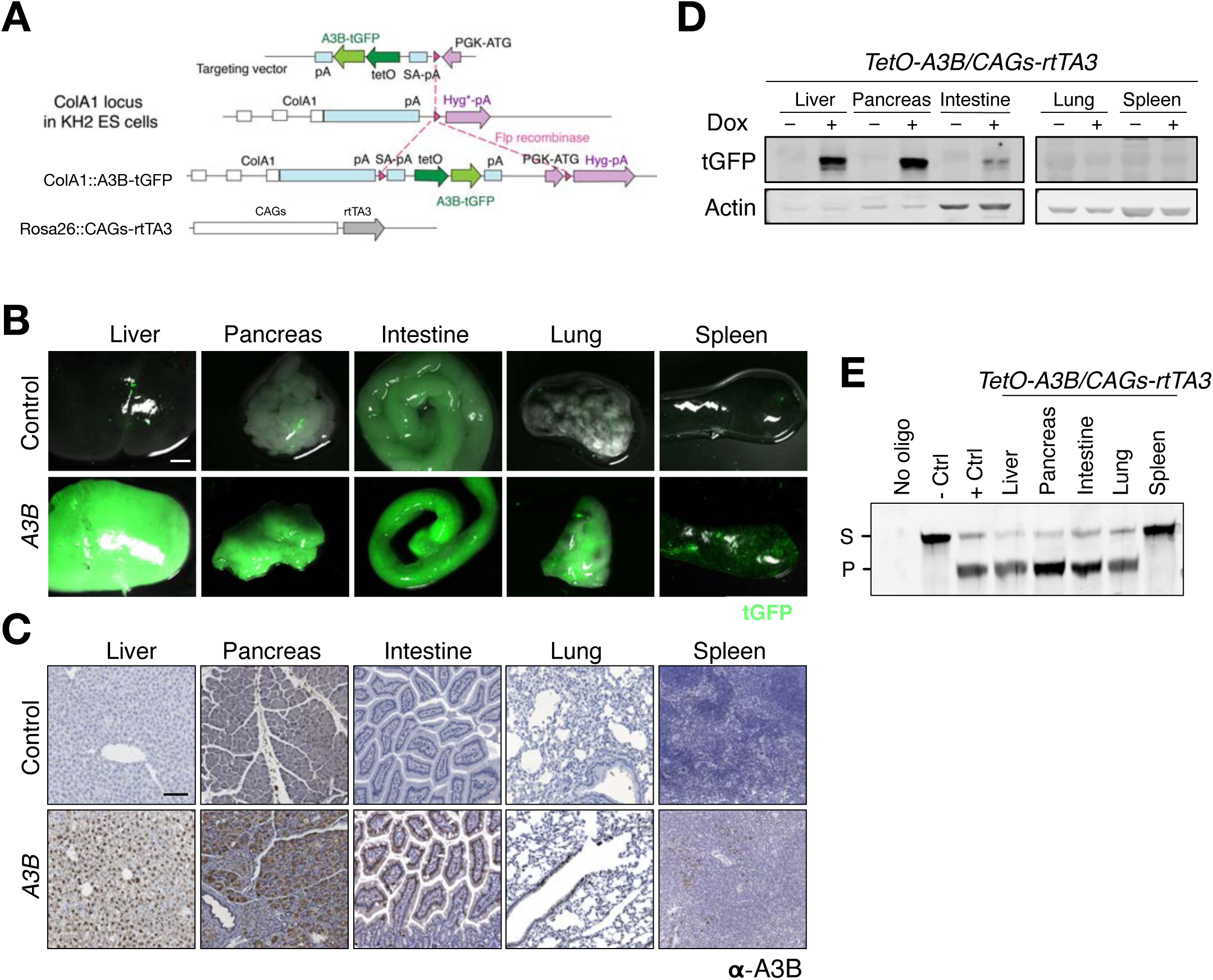
Generation of A3B-inducible mice. **A**) Schematic representation of the strategy used for the generation of human A3B transgenic mice. Under a TRE promoter, the human *A3B* cDNA fused to *tGFP* was inserted after homologous recombination into the *ColA1* locus of KH2 ES cells. **B**) Macroscopic images demonstrate tGFP fluorescence in tissues from *TetO-A3B/CAGs-rtTA3* mice fed with doxycycline for 10 days (scale bar: 3 mm). **C)** Immunohistochemistry of A3B in the indicated tissues from *TetO-A3B/CAGs-rtTA3* mice fed with doxycycline for 10 days (scale bar: 100 µm). **D**) Western blot analysis showing A3B levels in the indicated tissues from *TetO-A3B/CAGs- rtTA3* mice with and without doxycycline treatment for 10 days. Anti-actin blots are shown as loading controls. **E)** DNA cytosine deaminase activity from whole cell extracts in the indicated tissues (S, Substrate; P, Product).

### Acute APOBEC3B induction *in vivo* is lethal

To examine the consequences of expressing A3B *in vivo*, dox-containing food was given to adult *TetO-A3B/CAGs-rtTA3* mice. Unexpectedly, *A3B* mice showed a rapid health deterioration, and all animals died within 6 to 14 days after dox administration, while control animals were healthy beyond 365 days (Figure 2A). Prior to death, A3B expressing mice were largely unresponsive and immobile with a clear ‘‘trembling’’ phenotype. Macroscopically, *A3B* livers appeared indicative of fatty acid accumulation and close examination revealed micro-and macro-vesicular steatosis (Figure 2B). Histopathological characterization suggested loss of tissue architecture and abnormalities in several organs incompatible with life. Differential expression analysis of RNA-sequencing (RNA-seq) data comparing control and *A3B* expressing livers indicated a metabolic disturbance, due to the downregulation in the cholesterol metabolism (Figure S2A). Analysis of paraffin sections revealed an increase in apoptosis and DNA damage (Caspase3 and γH2AX staining, respectively) in *A3B* livers compared to controls (Figure 2C), whereas no differences in proliferation were observed (Figure S2B). Increased serum levels of liver enzymes, such as alanine transaminase (ALT) and aspartate transaminase (AST), also indicated liver damage (Figure S2C).

**Figure 2.**
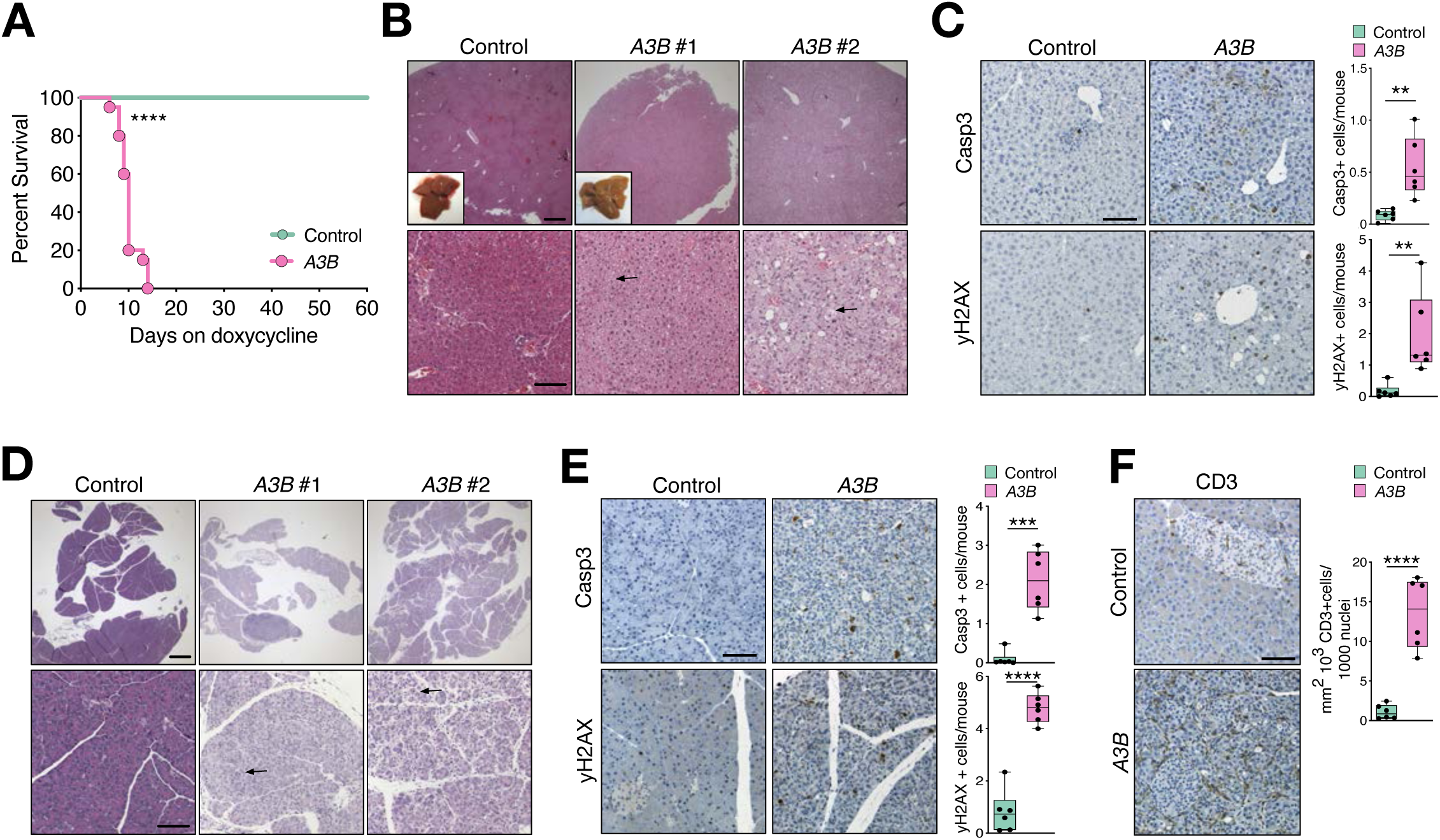
A3B overexpression causes early lethality. **A**) Survival of *TetO-A3B/CAGs-rtTA3* mice after doxycycline administration (controls n= 11; A3B n=12; P<0.001 by Log-rank (Mantel-Cox) test). **B)** H&E stained sections of livers from control and *A3B* mice (insets: macroscopic images). Arrowheads point to regions showing micro or macro-vesicular steatosis. **C)** Immunohistochemistry against Caspase3 and ψ−H2AX in liver sections from control and *A3B* mice and corresponding quantification (n=6). **D)** H&E stained sections of the pancreas from control and *A3B* mice. Arrowheads point to regions showing acinar destruction. **E)** Immunohistochemistry against Caspase3 and ψ−H2AX in paraffin sections of pancreas from control and *A3B* mice and corresponding quantification (n=6). **E)** Immunohistochemistry for CD3 in paraffin sections of pancreas from control and *A3B* mice and corresponding quantification (n=6). Data in panels C, E and F were analyzed by unpaired t test **p < 0.01, ***p < 0.001, ****p < 0.0001. Data represented as mean ± SD shown by dots, where each dot represents a mouse, and error bars, respectively. Scale bars: 100 µm. Scale bars upper panels B and D: 500 µm.

Parallel analyses of pancreatic sections, which also presented high levels of A3B, revealed signs of chronic pancreatitis and subsequent reactive hyperplasia leading to the disruption of the tissue architecture and destruction of the acinar cells (Figure 2D). Mild to moderate chronic inflammation, as well as increased number of neutrophils, were detected in the surrounding mesenterial adipose tissue. Reactive hypertrophy of regional lymph nodes was also observed. These findings correlated with upregulation of inflammatory response pathways identified by differential expression analysis in the *A3B* pancreas compared to control (Figure S2D). Increased apoptosis, DNA damage and T lymphocyte infiltration were also observed by immunohistochemistry in *A3B* pancreas (Figure 2E and 2F). Similar to liver tissues, no differences in proliferation were found (Figure S2E). Altogether, these results suggest that high levels of A3B lead to prohibited cell dysfunction and consequently disruption of tissue homeostasis, systemic organ failure, and early animal death.

### RNA editing by APOBEC3B

A3B has been implicated in the generation of DNA damage, mutagenesis, larger-scale chromosomal instability and phenotypic heterogeneity in cancer, mainly through its single-stranded DNA deamination activity (Burns, Lackey, et al., 2013; Hoopes et al., 2016). The related A3A enzyme, which has high homology with the catalytic domain of A3B (92% identity), has been shown to be capable of editing RNA cytosines in primary human cells (macrophages) and human tumors (Jalili et al., 2020; Sharma et al., 2015) further implicating this enzyme in tumor mutation and evolution (Cortez et al., 2019; Isozaki et al., 2021; Law et al., 2020). We therefore explored whether A3B could function as an RNA editing enzyme *in vivo*.

To test A3B RNA editing activity, RNA-seq and whole exome sequencing (WES) were performed in liver and pancreatic tissues (high A3B levels) from *A3B* mice after dox administration (10-14 days) as well as from similarly aged control littermates. After applying quality filters and performing data analyses, all single base changes at the DNA or RNA level were considered as A3B-specific changes if represented by >5% altered reads in *A3B* samples and no altered reads in control samples. Changes that were not observed from the matched exome data for each sample were used for RNA editing analysis (Figure 3A). A3B expressing livers and pancreas (6 samples each) presented a high number of edits with an average of 158 +/ 47.8 in liver and 285 +/ 56.2 in pancreas. Analysis of the different types of editing in the liver showed a global increase in A-to-G changes, potentially due to endogenous ADAR enzymes, as well as C-to-U changes (Figure 3B and S3A). Similarly, in the pancreas, even though the majority of the changes included A-to-G changes an elevated contribution of C-to-U changes was also detected (Figure 3C and S3B). Although an increased number of changes at the DNA level were also found, changes at A3B-preferred TCW motifs were a minority. This result was not surprising since the early death of the mice did not allow for the generation of clonal mutations to be detected by bulk DNA sequencing (Figure S3C and S3D).

**Figure 3.**
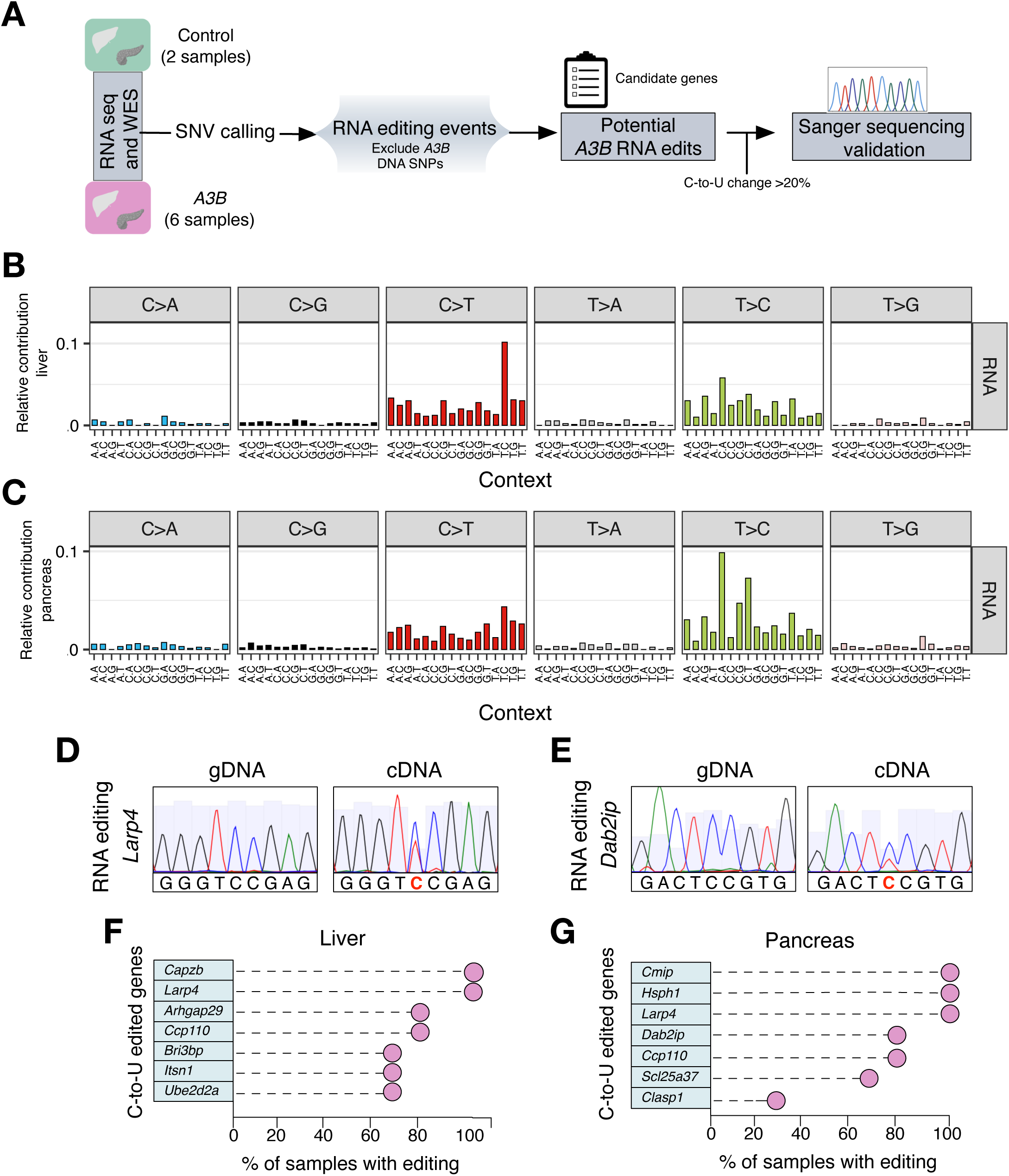
Local preference of APOBEC3B-driven RNA editing. **A**) Schematic of the pipeline used to call RNA-editing sites in liver and pancreas. **B-C**) Trinucleotide mutation profiles for all base substitutions in RNA from liver and pancreatic tissues of *A3B* mice (n=6 in each group). **D-E**) Examples of Sanger sequencing chromatograms showing A3B-driven RNA editing in *A3B* liver and pancreatic tissues, respectively. **F-G**) Lollipop plots of RT-PCR experimental validation of the indicated C-to-U editing targets with editing percentages shown (n=6 tissues in each group).

From the total RNA edits, we randomly selected potential candidate genes (C-to-U >20%) which were further validated to corroborate the reliability of our analysis (Figure 3A). We identified 7 positions that showed C-to-U changes in liver and another 7 positions in pancreas for experimental validation of site-specific editing by Sanger sequencing of purified DNA and RT-PCR RNA products (Figure 3D and 3E). In addition, to further explore whether the RNA editing events found in the *A3B* liver samples were random or recurrent events due to A3B overexpression, we sequenced these 7 edited positions in 6 additional *A3B* expressing mice. In the liver, we found that 2 of these positions were edited in 100% of the *A3B* mice, 2 positions were edited in 5 out of 6 mice (83%), and the last 3 positions tested were edited in 4 mice (66%) (Figure 3F). In the pancreas we found similar results, with 3 of the validated positions edited in 100% of the *A3B* mice, 2 positions edited in 5 out of 6 mice (83%), 3 positions in 4 out of 6 mice (66%), and the last position in 2 out of 6 mice (33%) (Figure 3G). Altogether these results suggest that certain sites may be hotspots for A3B editing.

To explore whether moderate levels of A3B found in other tissues such as lung (Figure S3E) also induced RNA editing, we performed RNA-seq of 6 controls and 6 *A3B* lung tissues. After data processing and analysis, we generated a list of potential candidate genes and experimental validation by Sanger sequencing of candidate C-to-U changes identified them as DNA SNPs (Figure S3F), suggesting that moderate levels of A3B do not induce RNA editing at least at detectable levels by Sanger sequencing.

### APOBEC3B-driven RNA editing occurs at a specific hotspot

The majority of the A3B induced RNA changes were detected in a UCC context and all the edited genes that were validated in liver and pancreatic samples were present in at least two animals. Therefore, we next explored whether A3B has any nucleotide preference surrounding the edited sequence. Analysis of the flanking nucleotides identified a broader nucleotide context, 5’-U<u>C</u>CGUGUG, surrounding the edited cytosine which could function as a predictor of A3B catalyzed RNA editing sites in the liver and pancreas of these mice (Figure 4A). In addition, no predicted stem-loop structures with the reactive C at the 3′-end of the loop were found in these regions in contrast to prior reports for A3A RNA editing hotspots (Jalili et al., 2020; Sharma & Baysal, 2017). The majority of the C-to-U modifications occurred at 3’UTRs, while 30% of the C-to-U sites were in coding exons, causing 1 stop, 13 non-synonymous and 23 synonymous changes (Figure 4B). Extensive analysis from the RNA edited positions revealed that 46 edited sites were recurrent in the liver from different samples, while 71 sites in the pancreas occurred across different mice (Figure S4A and S4B). More interestingly, we found that 65 positions were shared between the liver and the pancreas (Figure S4C) (Table 1). Altogether, these results reinforce our finding that A3B has RNA editing activity in a UCC- specific context which could be interpreted as hotspots for A3B-driven RNA editing.

**Figure 4.**
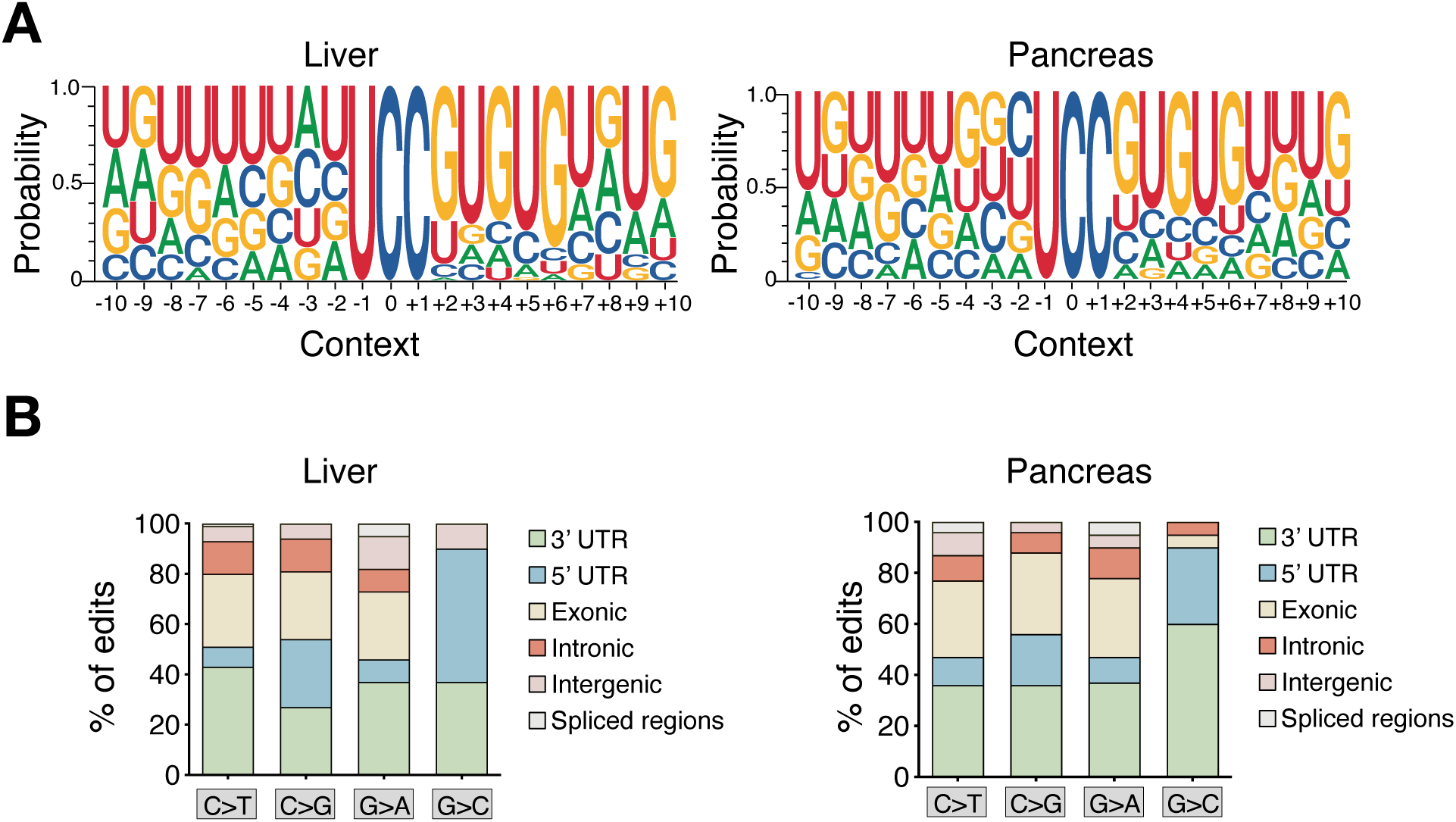
APOBEC3B-driven RNA editing occurs at a specific hotspot and mainly at 3’ UTRs. **A**) Web logo representations of the broader sequence preferences surrounding the C-to-U editing events in 5’-UCC motifs in liver (left) and pancreas (right). **B**) Distributions of the editing sites by type of RNA editing in liver (left) and pancreas (right).

### Endogenous APOBEC enzymes are not responsible for the C-to-U edits in *A3B* mice

The APOBEC family members in rodents include *Apobec1, Apobec2, Apobec3*, and *Aicda* (AID). To address, whether these other enzymes could be responsible for the editing events and the phenotype observed, we assessed the endogenous expression levels of the different *Apobec* and *Adar* genes in liver specimens using RNA-seq data. Expression levels of these genes (*Aicda*, *Apobec1*, *Apobec2*, *Apobec3*, *Adar*, *Adarb1*, and *Adarb2*) were similar between controls and A3B overexpressing samples, making it unlikely that the RNA editing events were due to the activity of one of these enzymes (Figure 5A).

**Figure 5.**
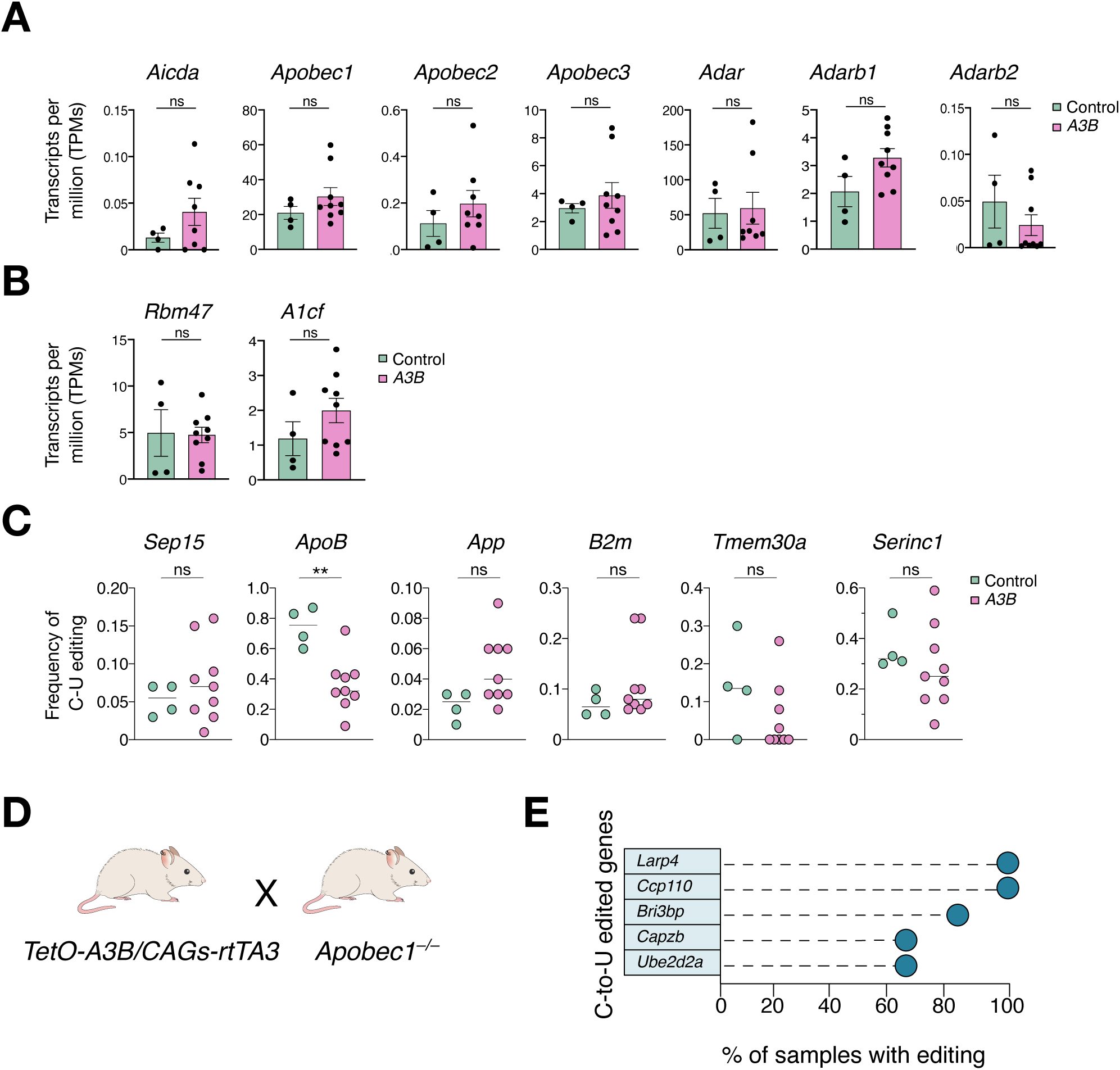
Endogenous Apobec enzymes are not responsible for the observed RNA editing. **A**) Average expression levels from the different endogenous *Apobec* and *Adar* family members obtained from all RNAseq data (transcripts per million (TPMs); each dot represents data from one animal n=4 controls and 9 *A3B* livers). **B**) Average expression levels from *Apobec1* cofactors obtained from the RNA sequencing data and showed as transcripts per million (TPMs); each dot represents data from one animal (n=4 controls and 9 *A3B* livers). **C**) Frequency of C to U editing of mRNA *Apobec1* well-known editing sites measured by quantification of RNA-seq data from controls and A3B livers. Each dot represents data from one animal (n=4 controls and 9 *A3B* livers). **D**) Schematic of the breeding strategy to obtain *A3B/Apobec1^-/-^* mice. **E**) Lollipop plots of RT-PCR experimental validation of the indicated C-to-U editing targets with editing percentages shown in *A3B/Apobec1^-/-^* mice (n=6 liver tissues).

APOBEC1 is an established RNA editing enzyme in the mouse liver (Blanc et al., 2014; Rosenberg et al., 2011; Teng et al., 1993). We then evaluated whether A3B overexpression might interfere with endogenous APOBEC1 functionality and cause the observed editing. To address this point, we examined the expression of the *Apobec1* cofactors required for editing. We found that neither *Rbm47* nor *A1cf* expression were changed in *A3B* livers compared to controls (Figure 5B). Next, we checked well-known RNA editing sites for *Apobec1* (Blanc et al., 2014) and found no significant differences in the editing frequencies from the majority of the *Apobec1* targets, except for the *ApoB* transcript, where editing was reduced by half in *A3B* livers compared to controls (Figure 5C). Consequently, since *Apobec1* showed reduced editing activity in its main target ApoB, the newly detected RNA editing events were unlikely an effect of *Apobec1* deamination but more likely a direct consequence of A3B overexpression. To further address this point, we crossed *A3B* mice with *Apobec1* knockout animals (Hirano et al., 1996) (Figure 5D). Shortly after doxycycline administration (6-11 days) triple transgenic mice succumbed to death, suggesting that the detrimental effect of A3B is independent of murine *Apobec1* (Figure S5A). *A3B/Apobec1^-/-^* mice express A3B at high levels comparable to single *A3B* animals and the deaminase activity of A3B was not altered due to *Apobec1* loss (Figure S5B). In addition, several of the recurrent RNA editing events described above were still evident in the *A3B/Apobec1^-/-^* livers (Figure 5E). Altogether, these data show that A3B and not endogenous *Apobec1* is responsible for the observed RNA editing events.

### Continuous APOBEC3B expression is required for RNA editing

To understand whether continuous expression of A3B is required to edit the RNA, we took advantage of the doxycycline-inducible model which allows to silence the transgene expression after dox withdrawal. Mice were fed with doxycycline for 4 days, followed by doxycycline deprivation to abrogate *A3B* expression and liver samples were collected at defined time points. First, we analyzed liver samples 12 days after doxycycline withdrawal. Second, mice received normal food until the end of the experiment (1 year). Third, mice expressed A3B in cycles (4 days on dox plus 26 days off dox) every month until the end of the experiment (1 year) (Figure 6A). We observed that 4 days of doxycycline administration were sufficient to trigger *A3B* expression and consequently induce A3B-driven RNA editing events (Figure 6B). Moreover, doxycycline withdrawal for 12 days was enough to abrogate A3B expression and subsequently A3B-driven RNA edited positions could no longer be detected (Figure 6C). Accordingly, mice expressing A3B in a pulse or a cycle manner showed no expression of A3B and no detectable RNA editing (Figure 6D-E). Altogether, these results indicate that high and continuous expression of *A3B* is required for RNA editing.

**Figure 6.**
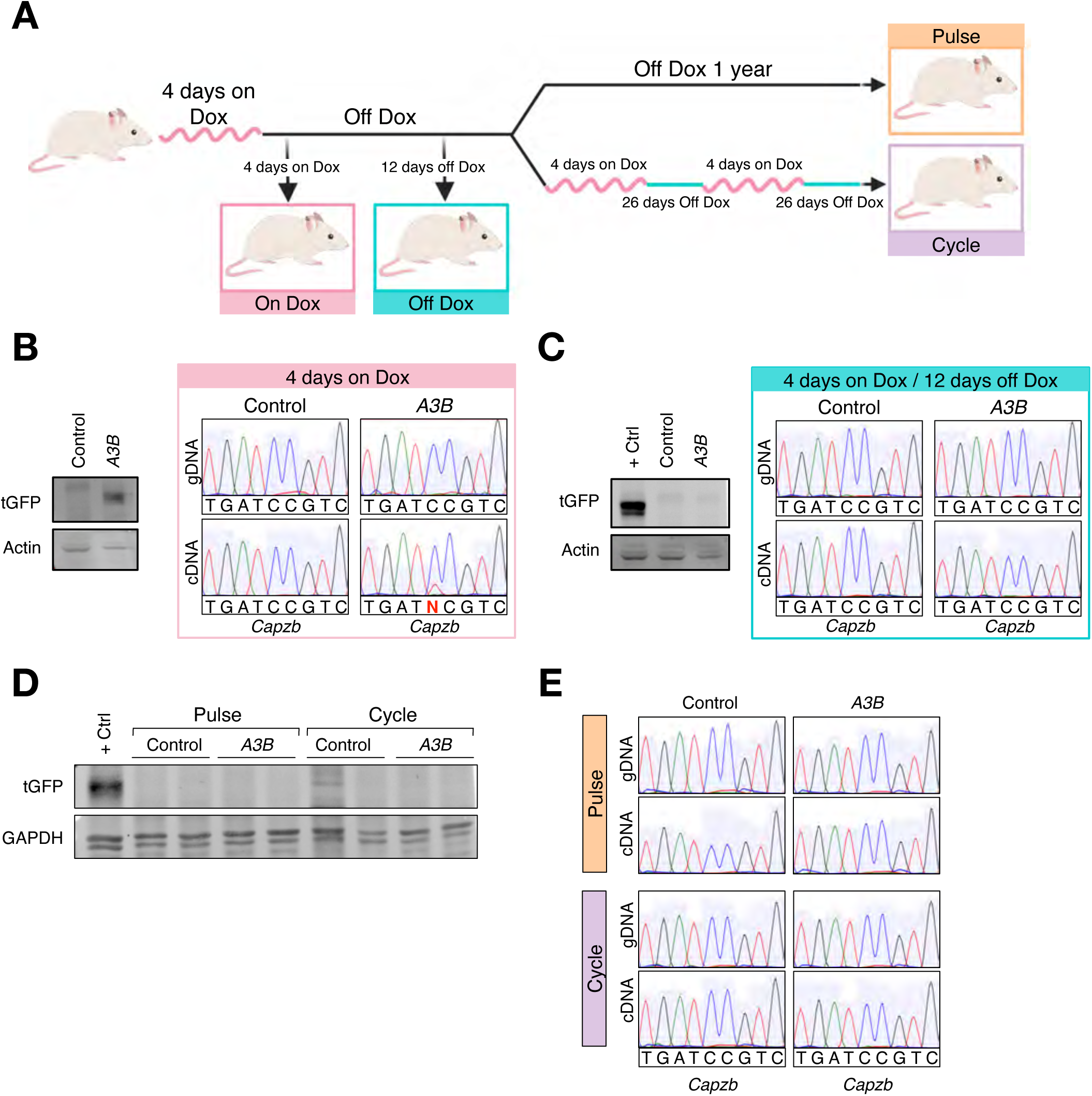
Continuous expression of A3B is required for RNA editing. **A)** Schematic representation of the strategy used to study whether A3B expression is needed to detect the RNA edits**. B), C)** and **D**) Western blots showing A3B levels in liver tissues from *A3B/CAGs-rtTA3*. Anti-actin or Anti-GAPDH is shown as the loading control. Example of Sanger sequencing chromatograms from the same livers for the A3B-driven edited positions. **B)** Mice 4 days after doxycycline administration. **C**) Mice on dox for 4 days and placed back on normal diet for 12 days. **D**) Mice that received a pulse of A3B expression (4 days dox and up to a year on normal diet) or expressing A3B in a cycle manner (4 days dox-26 days off dox/monthly). Samples were collected at experimental end point (1 year). **E**) Examples of Sanger sequencing chromatograms showing A3B-driven RNA editing in *A3B* liver from mice under the pulse and cycle experiments.

## Discussion

A3B, a single stranded-DNA mutagenic enzyme, is in the spotlight of cancer research as a driver of tumor evolution and therapeutic resistance. Several studies have attributed A3B- induced tumor heterogeneity to its mutagenic activity (Burns, Lackey, et al., 2013; Henderson et al., 2014; Leonard et al., 2013; McGranahan et al., 2015). Moreover, a recent study has shown that A3B-induced DNA damage can also contribute to chromosomal instability and increased tumor heterogeneity (Venkatesan et al., 2021). However, whether A3B can also influence tumorigenesis by RNA editing activity remains unknown. Employing a novel doxycycline inducible mouse model, in which high and sustained A3B levels were achieved, we found that A3B is not well tolerated by the cells and compromises animal survival by inducing DNA damage and inflammation. Importantly, we show that A3B catalyzes C-to-U deamination in the RNA. A3B was previously described to solely deaminate genomic DNA showing C-to-U activity of the enzyme on ssDNA but until now studies have failed to show RNA C-to-U editing activity in any model system. This is therefore the first demonstration *in vivo* that human A3B has the capacity to edit cytosines in RNA.

Several APOBEC members have a dual function in editing RNA and inducing mutations in DNA (Pecori et al., 2022). However, the molecular basis that determines substrate range and selectivity warrants further investigation. Our analyses show that the most frequently edited trinucleotide in RNA in *A3B* expressing mice is UCC-to-UUC. In contrast, A3B- catalyzed mutations in DNA occur predominantly in a TCA and TCT motifs and TCC is strongly disfavored. An atomic explanation for this difference is not obvious, in part because only A3-ssDNA complexes have been determined by x-ray crystallography (Shi et al., 2017). However, one clue may derive from our observation that A3B-dependent RNA edits also have a broader sequence motif U<u>C</u>CGUGUG. Similarly, APOBEC1 edited RNA cytosines also appear to be part of a broader sequence context (AU-rich), and A3A deaminates RNA cytosines in the loop region of stem-loop structures (Jalili et al., 2020; Lerner et al., 2018; Pecori et al., 2022). These observations suggest that each enzyme has unique substrate preferences that may be related to their underlying biological functionality. Moreover, the analysis of the sequence context and secondary structures surrounding the edited cytosine could be used as a marker for activity by a particular deaminase family member (i.e., enzyme-specific biomarkers).

The RNA C-to-U editing activity of the APOBEC family is poorly understood. Few APOBEC members have been described to have RNA editing activity, highlighting the importance of considering other A3s for a possible role in editing. One of them, A3A which shows high homology and is evolutionarily related to A3B has RNA editing activity (Sharma et al., 2015), supporting the idea that A3B could have similar characteristics and also function as an RNA editing enzyme. Notably, A3A is biochemically more active than A3B, which could explain why the majority of events have been ascribed to A3A (Byeon et al., 2016). Moreover, A3A editing has been described to occur at hotspots in RNA stem-loops in human tumors expressing high levels of A3A (Jalili et al., 2020). Taking advantage of our doxycycline inducible mouse model we demonstrate that high A3B expression is required for editing, as control (non-A3B) animals elicited no editing and withdrawal of dox shuts-off RNA editing. In addition, our findings clearly indicate that A3B does not randomly deaminate cytosines and that there are hotspots for A3B-editing based on the sequence context but not on secondary structures. Therefore, focus on stem-loop structures for the identification of editing events and the possibility that high levels of A3B may be selected against in cancer cells, could have masked the A3B editing landscape. Additional studies in which the expression of A3B is monitored should focus on studying whether hotspots for A3B-editing may associate with (or even directly promote) cancer or other diseases.

Although Apobec1 and C-to-U editing have been linked to several biological processes, their dysregulation has also been associated with multiple diseases, including cancer (Luo et al., 2021; Swanton et al., 2015). Under these circumstances, the mechanisms regulating the correct enzyme activity might be lost and substrate availability might differ. For instance, Apobec1 overexpression in the liver of transgenic animals induces hepatocellular carcinoma, although whether this is the result of RNA editing or DNA mutation remains unclear (Yamanaka et al., 1995). Our model recapitulates A3B levels found in human cancer and generates a pathological environment. It is possible that when A3B is upregulated it deaminates DNA and RNA substrates indiscriminately and thereby compromises genome stability. We report here, for the first time that A3B, a known genome mutagenic enzyme, is also able to deaminate the RNA in mice, when overexpressed. We discovered that A3B-associated edits occur mainly at the UCC motif and that its expression is required to detect the edits. This evidence and the emerging collective findings in other family members suggest that the APOBEC family may utilize a combination of different mechanisms to induce genetic variability. These results highlight the importance of expanding our knowledge of C-to-U RNA-editing events. Because of the dynamic nature of RNA editing, it will be important to identify the timing when A3B is upregulated in human tumors to fully evaluate its implications.

## Supplementary Figure Legends

**Supplementary Figure 1.**
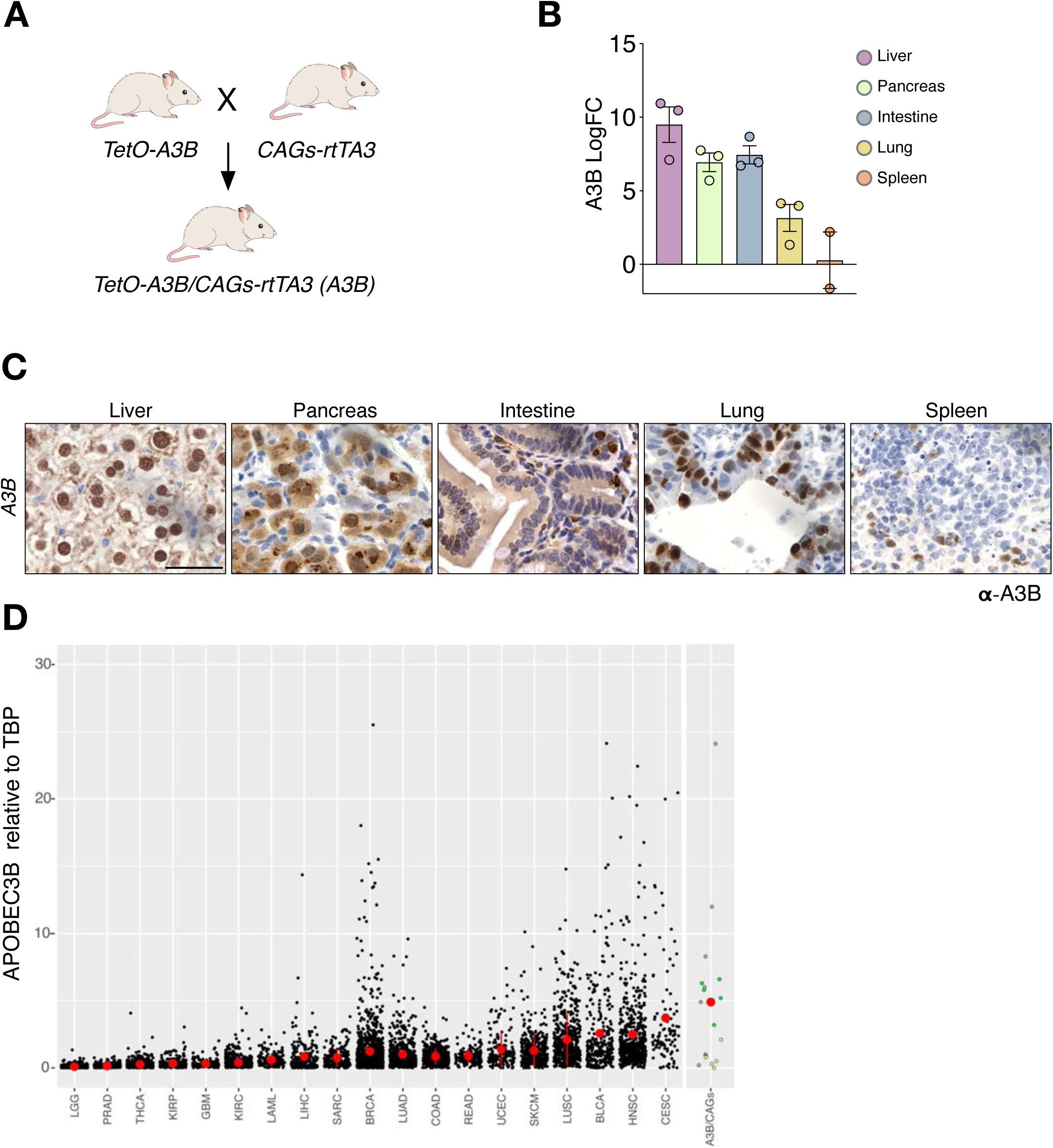
**A**) Schematic of the breeding strategy to obtain *TetO-A3B/CAGs-rtTA3* mice. **B**) Quantitative RT-PCR analysis of A3B transgene expression in different tissues from *A3B* mice after 10 days on doxycycline normalized to two housekeeping genes: Actin and 18S. Error bars represent SEM. **C**) Immunohistochemistry of A3B in the indicated tissues from *TetO-A3B/*CAGs-rtTA3 mice fed with doxycycline for 10 days. Scale bar: 50 µm. **D**) A3B mRNA levels in liver, lung and pancreas from *TetO-A3B/CAGs-rtTA3* animals are within the range reported for human tumors (gray dots represent data from individual TCGA tumors, purple dots represent liver, green dots pancreas and yellow dots lung tissues from *TetO-A3B/CAGs-rtTA3* mice; mean ± SD shown by red dots and error bars, respectively).

**Supplementary Figure 2.**
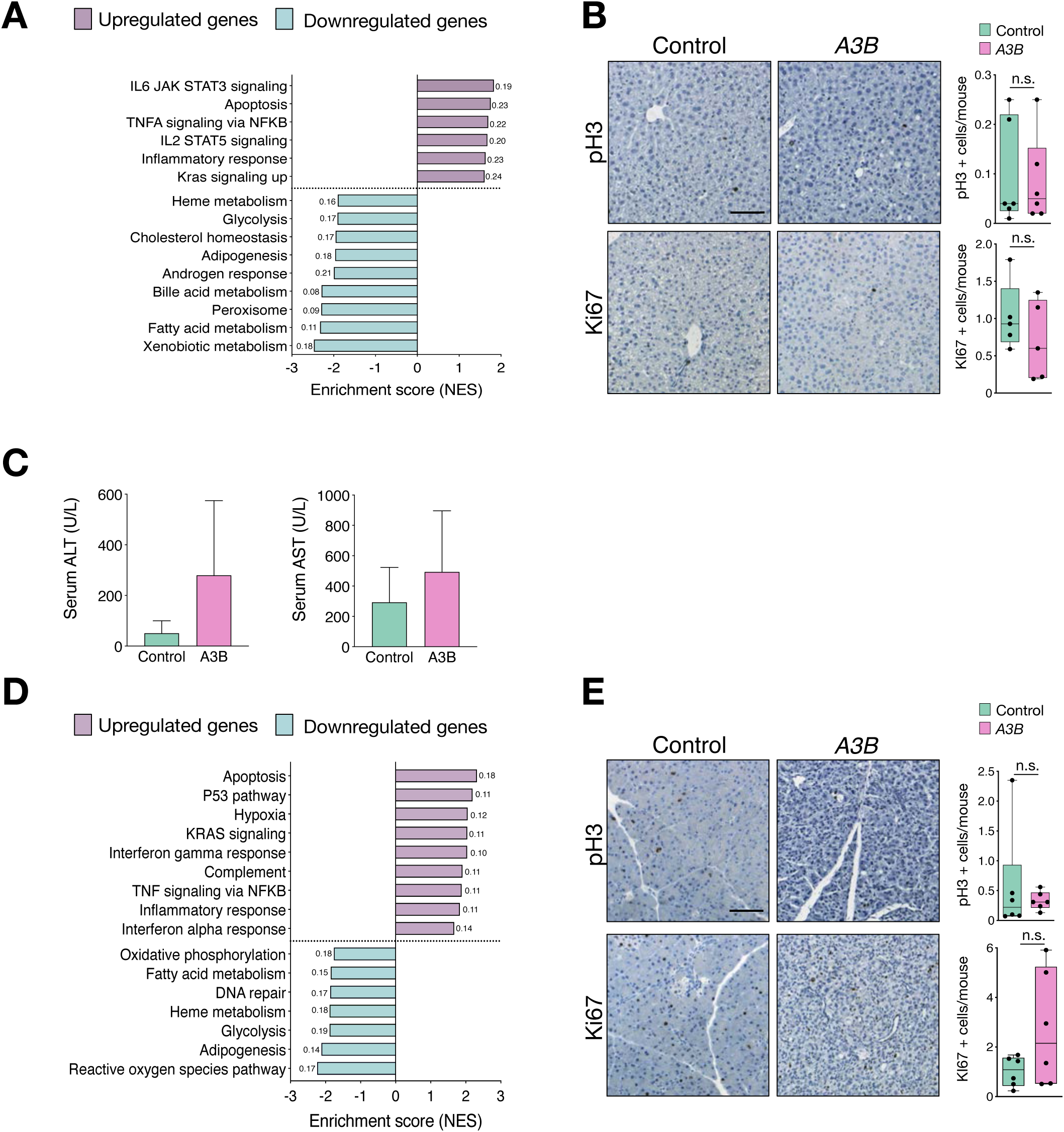
**A)** Pathway enrichment analysis showing the normalized enrichment score (NES) for the A3B liver samples compared to the controls. False discovery rate (FDR) q-values are shown to the right or left side of the bars. **B**) Immunohistochemistry against pH3 and ki67 in paraffin sections of livers from control and A3B mice and their corresponding quantification (n=6). **C**) Analysis of transaminase levels (ALT and AST) in sera of control and A3B mice. **D**) Pathway enrichment analysis showing the normalized enrichment score (NES) for the A3B pancreas samples compared to the controls. False discovery rate (FDR) q-values are shown to the right or left side of the bars. **E**) Immunohistochemistry against pH3 and ki67 in paraffin sections of pancreas from control and A3B mice and their corresponding quantification (n=6). Data was analyzed by unpaired t test, n.s. non-significant. Data represented as mean ± SD shown by dots and error bars, respectively. Scale bar: 100 µm.

**Supplementary Figure 3.**
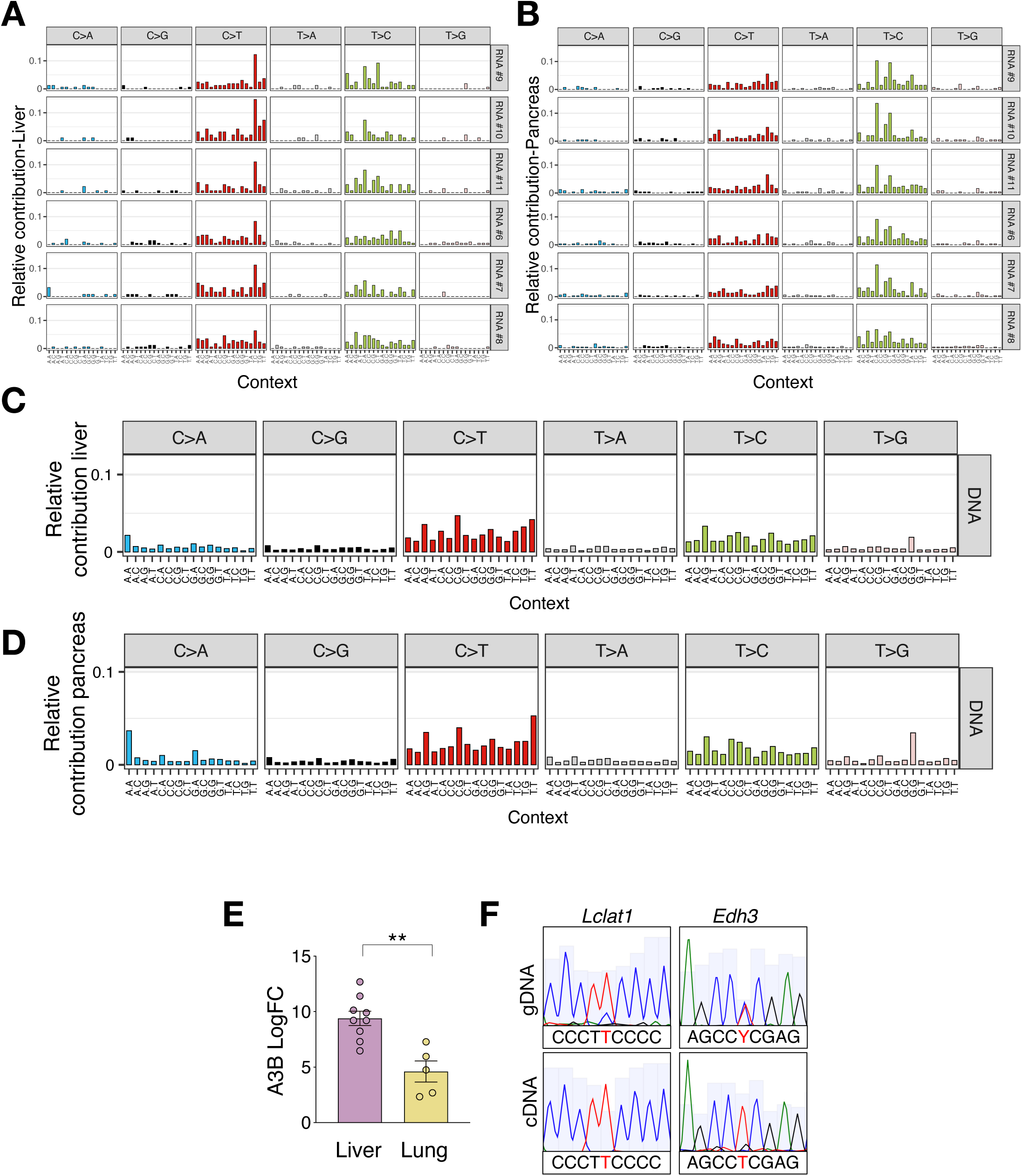
**A)** Trinucleotide mutation profiles for all base substitutions at the RNA level in livers from each individual A3B mouse. **B**) Trinucleotide mutation profiles for all base substitutions in pancreas from each individual A3B mouse. **C**) Trinucleotide mutation profiles for all base substitutions at the DNA level in livers from A3B mice, n=6. **D**) Trinucleotide mutation profiles for all base substitutions at the DNA level in pancreas from A3B mice, n=6. **E)** A3B mRNA levels in liver and lung from individual *A3B* mice relative to TBP (mean ± SEM shown by dots and error bars, respectively). **F**) Examples of Sanger sequencing chromatograms showing A3B- driven DNA mutations in A3B lungs.

**Supplementary Figure 4.**
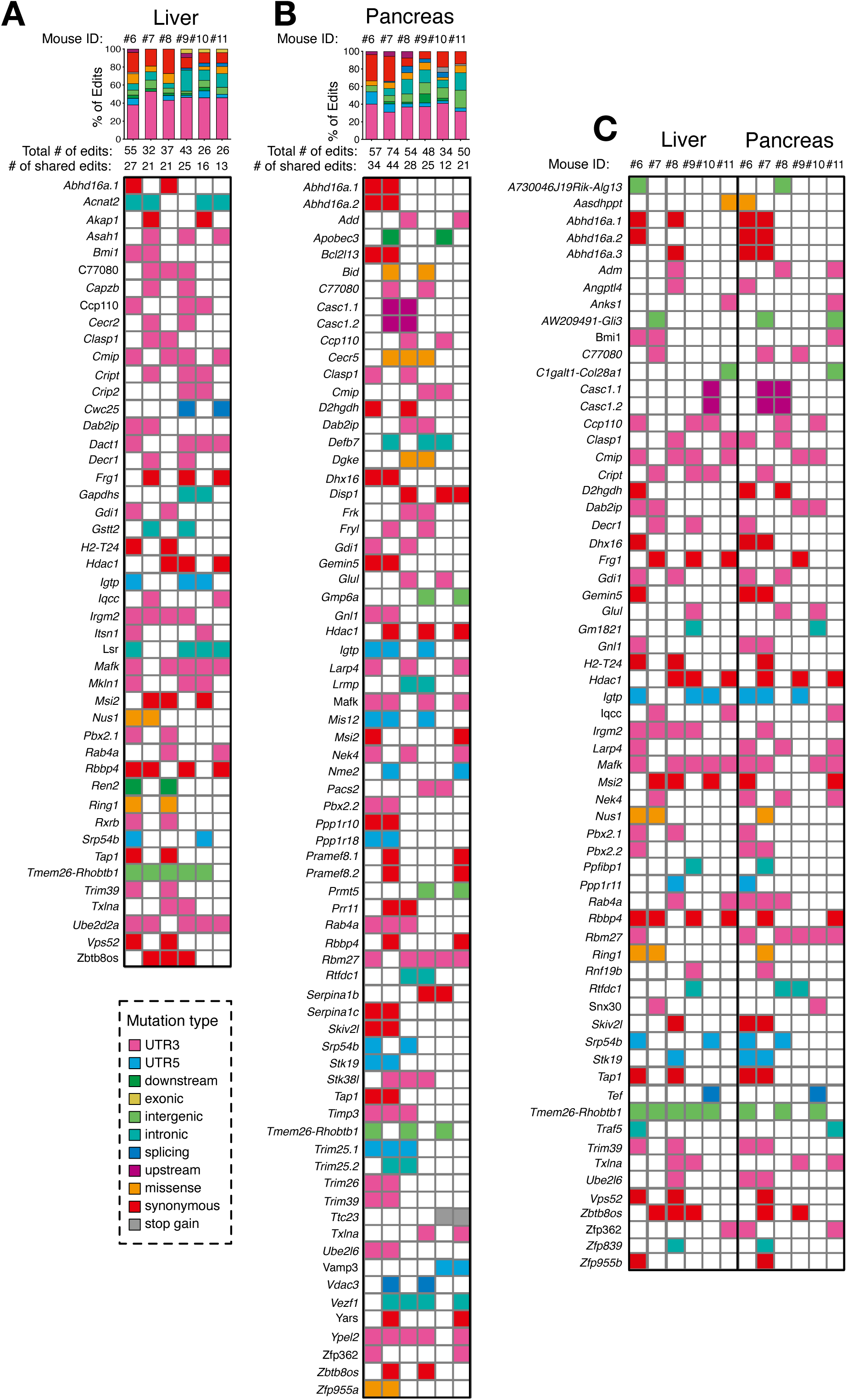
Editing landscape of shared edited genes in A3B-expressing livers and pancreas. The upper panel indicates the total number of edits per mouse and the lower panel the shared edits among mice. Represented only those edits that appeared in 2 or more mice. **A**) Liver **B**) Pancreas **C**) Liver and pancreas shared edits.

**Supplementary Figure 5.**
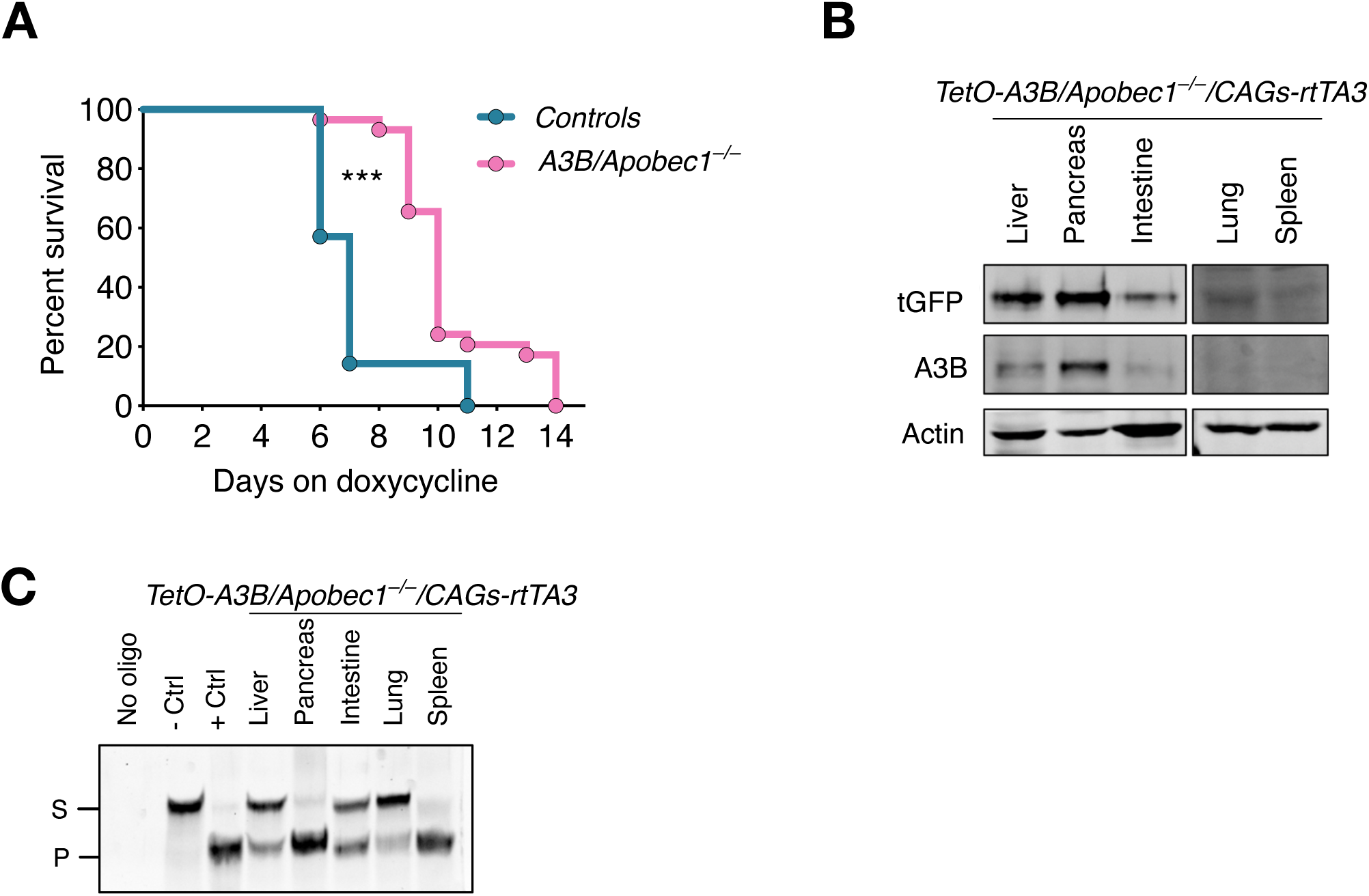
**A)** Survival of *A3B/CAGs-rtTA3* and *A3B/Apobec1-/-/CAGs-rtTA3* mice after doxycycline administration. *A3B* n= 29, *A3B/Apobec1-/-* n=7. P<0.0001 by Log-rank (Mantel-Cox) test. **B**) Immunodetection of tGFP in the indicated tissues from *A3B/Apobec1-/-* mice fed with doxycycline for 7 days. **C**) DNA cytosine deaminase activity from whole-cell extracts in the indicated tissues (S, substrate; P, product).

## Materials & Methods

### Mouse models

KH2 ES cells, were a gift from Sagrario Ortega and were generated by Konrad Hochedlinger and Rudolf Jaenisch (Beard et al., 2006). These ES cells carry the M2-rtTA gene inserted within the Rosa26 allele. A construct containing the human *APOBEC3B-tGFP* cDNA under the control of the tetracycline response element (TRE) was inserted downstream of the *Col1A1* locus. *ColA1-APOBEC3B*/*Rosa26-rtTA* heterozygous animals were bred out to exclude the *Rosa26-rtTA* transgene and bred to *CAGs-rtTA3* mice (Dow et al., 2014). All experiments were performed at the DKFZ animal facilities, with ethical approval from Baden-Wurttemberg, Germany (animal license No. G-29-19).

For inducing the transgene *in vivo*, different genotypes of the APOBEC3B:rtTA alleles [ (*+/A3B*)(*+/rtTA*); (+/+)(+/*rtTA*); (+/*A3B*)(*+/+*)] were fed with 625 ppm dox impregnated food pellets. For control, littermates with the genotypes (+/+)(+/*rtTA*); (+/*A3B*)(+/+) were fed with doxycycline.

Isolation of ear punch-DNA was performed via incubation in 100µL 0.05M NaOH at 98°C for 1h and subsequent neutralization with 10µL 1M Tris HCl ph7.5. *Rosa26-rtTA and CAGs-rtTA* transgenic mice were genotyped as described previously (Dow et al., 2014). The following oligonucleotides were used to genotype the *ColA1-A3B* allele: KH2-A3B A: 5’GCTGGGACACCTTTGTGTACCG 3’, KH2-A3B B: 5’ATCACGTGGCTCAGCAGGTAGG 3’. For all transgenes, the following PCR program was applied: 94°C for 2 min, 30 times [95°C for 30 s, 60°C for 30 s, 72°C for 30 s], and a final step at 72°C for 1 min.

### Immunodetection in tissue sections

Tissues were fixed in formalin overnight and embedded in paraffin. Antigen retrieval was performed using 0.09% (v/v) unmasking solution (Vector Labs, H-3300) for 30 minutes in a steamer. Inactivation of endogenous peroxidases was done using 3% Hydrogen Peroxide (Sigma, H1009) for 10 minutes. Secondary antibody staining and biotin-streptavidin incubation were performed using species-specific VECTASTAIN Elite ABC kits (Vector Labs, PK-6101 and BMK-2202). DAB Peroxidase Substrate kit (Vector Labs, SK-4100) was utilized for antibody detection. Primary antibodies used were anti-pH3 Ser10 (1:200, Cell Signalling, 9701), cleaved caspase 3 (1:200, Cell Signalling, 9661), yH2AX (1:200, Bethyl Labs 00059), APOBEC3B (1:200, a gift from R. Harris), tGFP (1:200 Origene, TA150041), ki67 (Medac 275R-18), CD3 (1:200, Dako A045229). Sections were visualized under a TissueFAXS slide scanning platform (TissueGnostics, Vienna, Austria). All the quantifications were done using StrataQuest software (TissueGnostics) to determine the number or percentage of pH3, Casp3, ki67, yH2AX and CD3 positive cells. Eosin G was from Roth and hematoxylin was from Linaris.

### Measurement of Serum Parameters

The analysis for AST, ALT was performed with mouse serum with the DRY-CHEM 500i analyzer (Fujifilm, Japan) following the manufacturer’s protocol.

### Deamination assay

DNA deaminase activity was measured in whole lysates of different tissues using previously described protocols (Law et al., 2016). A Fluor-labelled oligonucleotide containing a single target cytosine (5′-ATTATTATTATTCGAATGGATTTATTTATTTATTTATTTATTT-fluorescein) was incubated 3h at 37°C with the tissue lysates containing or not A3B. Samples were run in a 15% denaturing acrylamide gel and deamination activity was detected by fluorescence using iBright CL1500 imaging system (Thermo Fisher Scientific).

### Western Blot Assay

For protein extraction and immunoblotting, mouse tissues were lysed in RIPA lysis buffer (0.25 M Tris-HCl pH 6.8, 2.5% glycerol, 1% SDS, and 50 mM DTT). Samples were then boiled for 10 min and cleared by centrifugation. Protein expression was assessed by immunoblotting using 40-90 µg of the lysates and probed using the following specific antibodies: anti-APOBEC3B (1:1000, a gift from R. Harris); anti-tGFP (1:1000 Origene, TA150041) GAPDH (1:2000 Millipore, CB1001) and anti actin (1:5000 Sigma, A2066)

### DNA and RNA isolation

Snap frozen tissues were ground with a mortar and pestle on dry ice. For total RNA and genomic DNA extraction, AllPrep DNA/RNA Mini (Qiagen, 80204) was used and cDNA synthesis was done using the QuantiTect Reverse Transcription Kit (Qiagen, 205313) according to the manufacturer’s instructions.

### Real-time PCR

Quantification using real-time PCR was initiated using 10 ng of cDNA with SYBR Green PCR Master Mix (2×) (Applied Biosystems, 4364346) in a LightCycler II^®^ 480 (Roche) Relative mRNA levels were calculated according to the ΔCt or ΔΔCt relative quantification method and were normalized to the examined house-keeping genes (18S; Actin; TBP) levels.

### RNA-seq and WES data processing

Libraries were prepared using 1.2 ug total RNA with TruSeq Stranded kit for Illumina platforms. Libraries were sequenced with Illumina HiSeq 2000 v4 technology (125-nucleotide paired-end reads). Before and after trimming we evaluated the RNAseq quality with FastQC (https://www.bioinformatics.babraham.ac.uk/projects/fastqc/). Quality control, including per-base quality, duplication levels, and over-representative sequences, passed all the checkpoints. RNA-seq reads were aligned to mouse genome mm10 using STAR/2.7.1a with basic two pass mode for realigning splice junctions enabled. Picard tools (version 2.18.16) were then used to mark duplicate reads, split CIGAR reads with Ns at the splice junctions. Mutect2 from GATK (3.6) was used to call RNA edits relative to the pool of normal RNA-seq data from pancreas/liver/lung tissues of litter mates. Editing events that passed mutect2 internal filter with at least 6 reads supporting the edit and a minimum of 20 total reads at the editing site and a variant allele frequency greater than 0.05 were used for downstream analysis.

TPMs levels of various genes were extracted by generating the read-count matrix using the Bioconductor packages GenomicAlignments and GenomicFeatures in R. RNA editing levels at Apobec1 target sites were extracted from the raw tables of the aligned reads before applying any filters for further analysis.

To perform the differential expression analysis the sequenced reads were aligned to the mm10 reference genome using kallisto v0.46.1. Raw counts were normalized and differentially expressed genes (DEGs) were calculated using R with the DEseq2 package. Gene set enrichment analysis (GSEA) of differentially expressed genes was performed using ‘ClusterProfiler’ package of R and javaGSEA Desktop Application v2.2.2. Pathways with an FDR-P value ≤ 0.25 were chosen as significantly enriched.

Libraries were prepared using 0.5 ug total genomic DNA with Agilent Low Input Exom-Seq Mouse kit for Illumina platforms. Libraries were sequenced with Illumina HiSeq 2000 v4 technology (125-nucleotide paired-end reads). Whole exome data were aligned to the mouse genome (mm10) using SpeedSeq (PMID: 26258291). PCR duplicates were removed using Picard (version 2.18.16). Reads were locally realigned around Indels using GATK3 (version 3.6.0) tools. Single base substitutions and small InDels were called relative to the pool of normal tissues from litter mates using Mutect2. SBSs that passed the internal GATK3 filter with minimum 4 reads supporting each variant, minimum 20 total reads at each variant site and a variant allele frequency over 0.05 were used for comparison to RNA editing events. RNA editing events that were not observed from the matched exome data for each sample were used for signature analysis.

All sequences logos were generated and visualized using “ggseqlogo” package in R.

### Candidate validation

Candidates were selected following the criteria that 10% of the transcripts for a specific gene must contain a C to T change, to ensure the visualization of a double peak after sanger sequencing. Primers were designed for the chosen candidates and used to amplify the edited position in RNA and DNA from livers and lungs. PCR products were run in an agarose gel and later bands were isolated and DNA extracted using QIAquick Gel Extraction kit (Qiagen). The extracted DNA was then submitted for sanger sequencing (Microsynth). Sequences were aligned to the specific reference mouse gene using the online tool Benchling to finally determine the presence of an RNA editing event or a DNA mutation. RNA editing events were additionally verified using MultiEditR v1.0.8

### Statistical Analysis

Statistical analysis was carried out using Prism6 (GraphPad). p values were as follows: *p < 0.05, **p < 0.01, ***p < 0.001, ****p < 0.0001. The number of animals is represented with n.

## Acknowledgments

We thank the DKFZ light microscopy unit, the DKFZ mouse facility and the Genomics and Proteomics unit for excellent technical assistance. We wish to thank Alberto Diaz, Maria Ramos, Sara Chocarro, Pan Fan, Cameron Durfee and Bojana Stefanovska for their suggestions on the manuscript. We are grateful to Mirian Fernandez-Vaquero and Prof. Dr. Mathias Heikenwälder for their help with the serum analysis of liver enzymes and to Darjus Tschaharganeh for providing *CAGs-rtTA3* mice.

A.A.V is supported by a grant from the Deutsches Zentrum für Lungenforschung (DZL, German Center for Lung Research). This work was in part supported by the DZL, # 82DZL004A4) to R.S and a grant to RSH from the National Cancer Institute (P01-CA234228). RSH is the Margaret Harvey Schering Land Grant Chair for Cancer Research, a Distinguished University McKnight Professor, and an Investigator of the Howard Hughes Medical Institute.

## Contributions

A.A.V and R.S. designed the experiments. A.A.V. performed the experiments and analyzed the data. N.A.T., R.T., E.R and U.B.D conducted bioinformatic analyses. K.S. generated the construct to develop A3B mice. N.P. provided the Apobec1 null mice. A.S and T.P. performed the pathology analyses. A.A.V, R.H. and R.S. wrote the manuscript with help from all authors. R.S. supervised the study.

